# Anlotinib overcomes multiple drug resistant of the colorectal cancer cells via inactivating PI3K/AKT pathway

**DOI:** 10.1101/821801

**Authors:** Weilan Lan, Jinyan Zhao, Haixia Shang, Jun Peng, Wujin Chen, Jiumao Lin

**Author notes:** Contributed equally (Weilan Lan, Jinyan Zhao). **Correspondence to:** Jiumao Lin (MD), Academy of Integrative Medicine, Fujian University of Traditional Chinese Medicine, 1 Qiuyang Road, Minhou Shangjie, Fuzhou, Fujian 350122, China, Wujin Chen (MD), Oncology Department, Affiliated People’s Hospital of Fujian University of Traditional Chinese Medicine, Fuzhou 350004, Fujian, China.

## Abstract

**Background:** Anlotinib is a multi-tyrosine kinase inhibitor that has been reported to have activity against colorectal cancer. However, the functional mechanisms whereby anlotinib mediates against deadly drug-resistant colorectal cancer (CRC) has not been fully described-specifically, the potential mechanisms that inhibit proliferation and induce apoptosis remain largely unknown.

**Methods:** MTT assays were used to detect cell viability and calculate the resistance index. Colony formation was used to evaluate the proliferation of resistant cells. DAPI staining was used to detect cell apoptosis morphologically. Annexin V-FITC with PI staining was used to detect early and late-stage apoptosis of cells. Cell cycle distribution was determined by Flow cytometry. Transwell assays were performed to examine the ability of migration and invasion. Cyclin D1 Survivin, CDK4, Bcl-2, Bax and changes of PI3K/AKT pathway were detected by Western blotting. Compared as a single agent or combined with anlotinib or LY294002, PI3K inhibitor (LY294002) was used to verify whether it inhibited drug-resistant CRC cells by lowering PI3K/AKT.

**Results:** HCT-8/5-FU cells showed multiple drug resistance. Drug resistance index of 5-FU, ADM and DDP were 390.27, 2.55 and 4.57, respectively. Anlotinib was shown to inhibit cell viability on HCT-8/5-FU and HCT-8 cells for 24 h and 48 h in a dosage- and time-dependent pattern. Compared with 48 h, intervened with anlotinib (0 μM, 10 μM, 20 μM and 40 μM) for 24 h, the HCT-8/5-FU cells were sensitive to anlotinib, and their sensitivity was greater than that of the parent cell line (HCT-8) at 24 h. Further, anlotinib inhibited the number of cloned cells significantly and had a significant inhibitory effect on cell cycle, mainly by blocking G1 transferring to S phase. Moreover, anlotinib could down-regulate the expression of survivin, cyclin D1, CDK4, caspase-3, Bcl-2, MMP-2, vimentin, MMP-9, and N-cadherin, while up-regulating cleaved-caspase-3, Bax and E-cadherin. Anlotinib inhibited the activity of the PI3K/AKT pathway and induced apoptosis in HCT-8/5-FU cells. Using LY294002, a specific PI3K inhibitor, our experiment found anlotinib can inhibit drug-resistant CRC cells by reducing PI3K and p-AKT activity-induce apoptosis.

**Conclusions:** Anlotinib inhibited the proliferation, metastasis and induced apoptosis of HCT-8 / 5-FU cells; and the mechanism could be that anlotinib overcomes multiple drug resistant of the colorectal cancer cells via inactivating PI3K/AKT pathway.

## Introduction

Colorectal cancer is a very common cancer associated with high mortality worldwide, the death rate of which ranks third among all malignant tumors[1-4]. In China, the incidence of CRC is increasing with each passing year by an average of 4% to 5%[5]. In recent years, great advances have been made in therapeutic regimens for treatment of primary CRC, and the mid-survival period has significantly lengthened. Prognosis, however, still remains poor, the long-term overall survival rates of CRC having remained largely unchanged over the past two decades[6, 7].

Currently, the main treatment for CRC is surgery. However, some patients with advanced-stage or recurrent CRC are not suitable for surgery and, thus, alternatively treated with chemotherapy using 5-fluorouracil (5-FU), Cisplatin (DDP), doxorubicin (ADM), vincristine (VCR) and so on. However, these drugs encountering resistance is one of the prime causes of the failure of clinical treatment. Accordingly, it is essential and extraordinarily pressing to explore novel therapeutic substances.

Anlotinib is a novel small molecule, a multi-target tyrosine kinase inhibitor, that has been used to treat various types of cancers, including CRC, based on its known ability to block angiogenesis[8]. It has proven strong inhibitory activity on many target receptors, for example PDGFR, FGFR and VEGFR[9-11]. Recently, it has been shown that many tumor cells are sensitive to anlotinib[8]. However, the mechanism has not been clearly elucidated. Therefore, it is essential to comprehend the intrinsic mechanism of anlotinib in the treatment of MDR colorectal cancer in order to improve its efficacy in clinical therapeutic application.

## Materials and Methods

### Reagents

RPMI 1640 medium, 0.25% trypsin and fetal bovine serum (FBS) were purchased from Gibco (Carlsbad, CA, USA). RIPA cell lysis buffer (pierce, 89,900) was obtained from Thermo Fisher Scientific, Inc. (Waltham, MA, USA). DAPI (cat.no.192133), FITC-Annexin V/propidium iodide (PI) apoptosis assay kit (cat. no. KGA108) was purchased from Nanjing KeyGene Biotech Co., Ltd. (Nanjing, China). Antibodies to cyclin D1 (cat. no. 60186), survivin (cat. no. 10508), CDK4 (cat. no. 11026), Bax (cat. no. 50599), Bcl-2 (cat. no. 12789), caspase-3 (cat. no. 19677), PI3K (cat. no. 60225), AKT (cat. no. 10176), p-AKT (cat. no. 66444), β-actin (cat. no. 66009), MMP-2 (cat. no. 10373) and MMP-9 (cat. no. 10375) were purchased from Proteintech (Chicago, IL, USA). N-cadherin (cat. no. 14215s), E-cadherin (cat. no. 14472s) and vimentin (cat. no. 3390S) were supplied by Cell Signaling Technology, Inc. (Danvers, MA, USA). LY294002 (cat. no. 28749) was purchased from MCE (Shanghai, China).

### Anlotinib hydrochloride preparation

Anlotinib hydrochloride was purchased from CTTQ (Chia Tai Tian Qing) Lianyungang Pharmaceutical Company (Jiangsu, China). Dissolved 12 mg anlotinib hydrochloride in 2.5 mL DMSO to 10 mM as the original solution and divided them into seven tubules stored at −20°C for in vitro experiments. When used, diluted to the required concentration.

### Cell culture

Human cell lines of colorectal cancer (HCT-8/5-FU and HCT-8) were purchased from Nanjing KeyGene Biotech Co., Ltd. (Nanjing, China). All cells were cultured in RPMI 1640 medium (RPMI1640+10% (v/v) FBS + 100 U/mL penicillin+ 100 µg/mL streptomycin). HCT-8/5-FU cells were maintained in RPMI 1640 medium containing 15 μg 5-FU, then according to the experiment aim respectively diluted into different cell densities for cells to other experiments. HCT-8/5-FU cells used in the follow up experiments were cultured completely in RPMI 1640 without 5-FU.

### Validation of cell resistance

MTT assay was used to validate cell drug resistance. HCT-8/5-FU and HCT-8 cells during logarithmic phase were seeded with the density of 1 × 105/mL cells in 96-well culture plate, 100 μL/well, cultured in 37 °C and 5% CO_2_ incubator, when cells convergence degree of 50%∼60%, remove the original culture medium, and then added different drugs 5-FU (0-25600 μM), ADM (0-256 μM) and DDP (0-256 μM). After treatment, removed the medium, adding MTT (0.5 mg/ml) to each well (100 μL), cultured approximately 4 h, removed supernatant, put DMSO (100 μL) in the well. The A value of cell viability was detected at 570 nm. SPSS 21.0 was used to calculate IC50, and then IC50 was used to calculate the drug resistance index (RI) RI =IC50 (HCT-8/5-FU)/IC50 (HCT-8).

### Cell viability assay

HCT-8/5-FU cell preconditioning was the same as MDR validation. Added anlotinib hydrochloride of different concentrations (0 μM, 1 μM, 10 μM, 20 μM and 40 μM) for 24 h or 48 h. After treatment, removed the medium, added MTT (100 μL, 0.5 mg/mL) to each well. After culturing approximately 4 h, removed supernatant, added 100 μL DMSO to each well. The value of A was measured at 570 nm. Cell growth inhibition rate (%) = 1- (experimental A value/control A value) ×100%.

### Colony formation assay

HCT-8/5-FU cells were inoculated 4.0×105 in 6-well plates and cultured overnight in cell incubator at 37 °C. When the cell confluence reached 50% ∼ 60%, added respectively anlotinib hydrochloride of different concentrations (0 μM, 10 μM, 20 μM and 40 μM) to intervene for 24 h. Collected 1000 cells were inoculated in 6-well plate for 10 days. When the clone appeared in the hole of culture plate, stopped culturing and washed with PBS 3 times, then added 4% paraformaldehyde to fix for 10 min. Discarded 4% paraformaldehyde and cleaned with PBS, crystal violet staining for 15 min. The colonies were calculated manually in the three different views.

### Cell cycle assay

Cell intervention as above. At the end of treatment, using 70% precooling ethanol to immobilize the cells in 4°C overnight and washed 3 times (PBS), 500 μL RI/Rnase A reaction staining in darkness for 30 minutes, and determined in FL2 channel of flow cytometry.

### DAPI staining

Cell intervention as above. Added 1 mL 4% paraformaldehyde to fix for 10 min. Put 1 mL DAPI staining solution in the well and cultured in cell incubator. After staining, washed the cells (PBS) 3 times. Observed and took some photographs with microscope (200×).

### Early cell apoptosis assays

Cell intervention as above. Annexin-V/PI staining was used to detect the early apoptosis of cells. Cell treatment was the same as cell colony formation assay. Collected supernatant and the cells, suspended with 500 μL binding buffer. Subsequently, put 5 μL Annexin V/FITC and 5 μL PI in the cell, respectively, cultured in the darkness for 15 min and determined in the FLI channel by Flow cytometry.

### Cell migration and invasion assays

Cell intervention as above. After treatment for 24h, 5×104 cells of HCT-8/5-FU were seeded in upper chamber coated with Matrigel (for invasion assay) or without Matrigel (for migration assay) with RPMI-1640 (without FBS); meanwhile, added 700 μL RPMI1640 to the lower chamber and cultured in cell incubator at 37°C for 14 h (invasion test) or 14 h (migration test). Added 4% paraformaldehyde in the upper and lower chambers to fix for 15 min, and crystal violet staining for 15 min. Cells in the upper layer of Transwell’s inner were washed and wiped clean. Then the lower layer of Transwell’s outer chamber was photographed with an inverted microscope (200×).

### Western blot analysis

Cell intervention as above. HCT-8/5-FU cells were collected and washed with chilled PBS. Cellular proteins were lysed by radioimmunoassay (RIPA) buffer with inhibitor cocktail (Thermo Fisher Scientific, USA), and centrifuged at 14,000 rpm for 20 min at 4°C to obtain final supernatants. Used BCA protein assay kit (bovine serum albumin) as a standard (Thermo Fisher Scientific, USA) to detect concentration of protein. Heated the compound (protein and loading-buffer) at 100°C for 8 minutes. Protein extract (30 μg) was loaded in the (10% SDS-PAGE) well, then used electrophoresis to separate protein and wet film and transferred to membrane for Western blotting. Lipid-free milk (5%) was used to seal the membrane for 1 hour and wash it. Then, the membrane was incubated overnight with the first-level antibodies of beta-actin, caspases-3 Bax, Bcl-2, cyclin D1, CKD4, p21, E-cadherin, N-cadherin, MMP-2, MMP-9, vimentin, PI3K, p-Akt and Akt (1:1,000) at 4 °C. Washed 3 times (TBST, 10 min), then incubated with secondary antibodies (HRP-conjugated 1:5000) for 1 h. Finally, we use Image Lab (Bio-Rad Laboratories, Inc., Berkley, California, USA) to detect protein.

### Statistical analysis

The statistical analysis of our experiments was carried out on a minimum of three independent experiments. IBM SPSS Statistics 21 was used to conduct statistical analyses. Compared with control group, all data with P values of less than or equal to 0.05 were considered to have statistical significance.

## Results

### HCT-8/5-FU cells had multidrug resistance but were sensitive to anlotinib

MTT assay was used to validate cell multidrug resistance and to determine the viability of HCT-8 or HCT-8/5-FU cells. The results showed that HCT-8/5-FU had multidrug resistance to different chemotherapeutic drugs, with resistance indices (RIs) of 5-FU, ADM and DDP of 390.27, 2.55 and 4.57, respectively (Table 1). Further, anlotinib could significantly inhibit cell viability of HCT-8/5-FU and HCT-8 cells in a time-and dosage-dependent fashion. However, anlotinib was more sensitive to HCT-8/5-FU cells in 24 h (Figure 1).

**Table 1.**
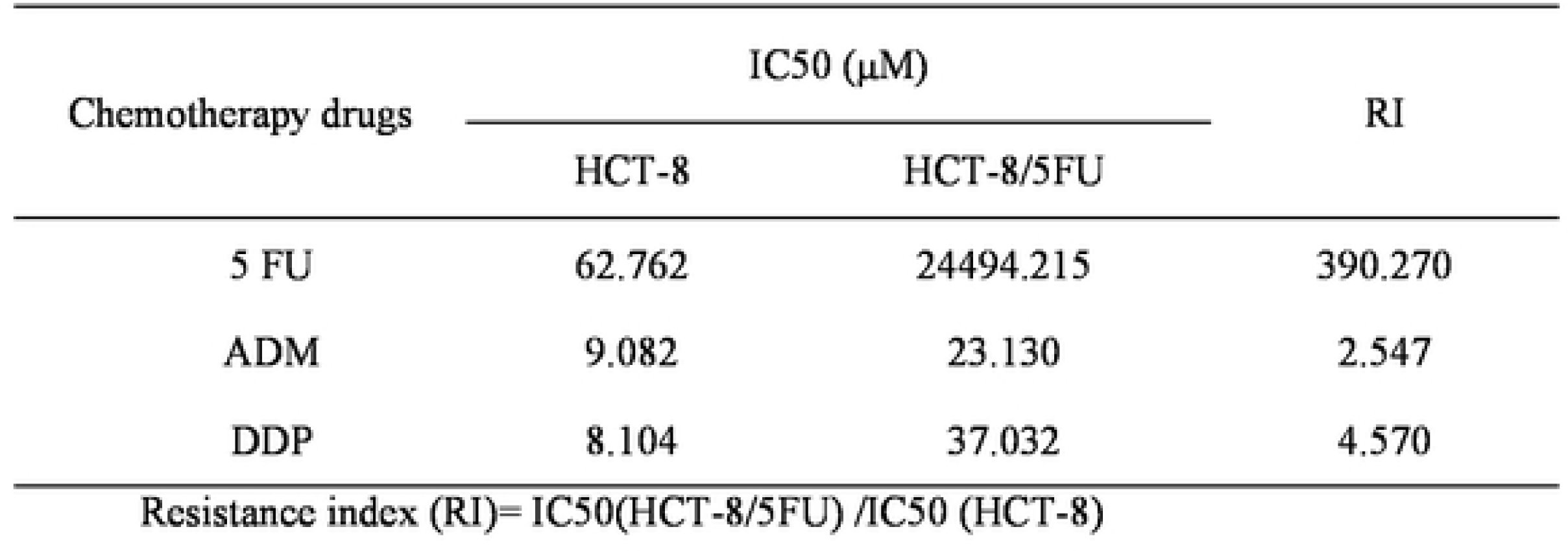
Resistance index of HCT-8/5-FU cells to different chemotherapeutic drugs. HCT-8 and HCT-8/5-FU cells were treated with 5FU, ADM and DDP for 48 h. Cell viability was tested by MTT assays. SPSS software was used to calculate the resistance index.

**Figure 1.**
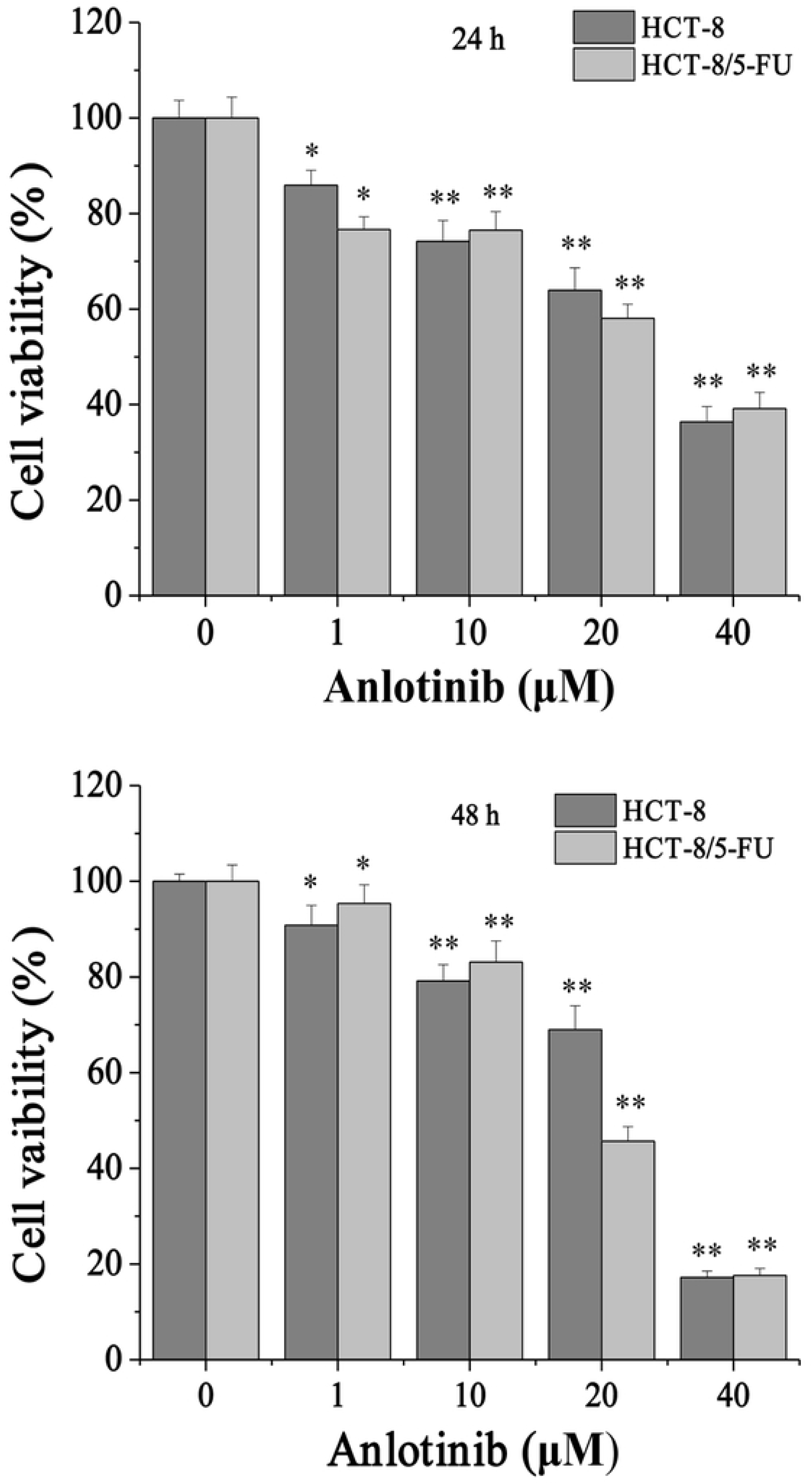
Anlotinib decreased cell viability of HCT-8/5-FU. HCT-8/5FU were treated with anlotinib (0, 1, 10, 20, 40 µM) for 24 h or 48 h. Then cell viability was calculated using MTT analysis. *P<0.05, **P<0.01.

### Anlotinib inhibited proliferation of HCT-8/5-FU cells via blocking cell cycles

To explore the mechanism of anlotinib activity in sensitizing multidrug-resistant HCT-8/5-FU cells, colony formation was performed. As shown in Figure 2A, compared to untreated control 587 ± 10, treatment with anlotinib (10, 20 and 40 μM) significantly decreased the cloned number (300 ± 11, 215 ± 10, 129 ± 11 respectively) (P<0.01). We next examined its influence in the HCT-8/5-FU cell cycle using FACS analysis with PI staining. As shown in Figure 2B, the percentage proportions of S-phase cells following treatment with 0, 10, 20 and 40 μM of anlotinib were 36.33 ± 0.85%; 11.74 ± 1.87%; 10.97 ± 1.32% and 7.89 ± 0.65 %, respectively (P<0.01). It is suggested that the inhibitory effect of Anlotinib on the proliferation of HCT-8/5-FU cells passes through G1/S cell cycle arrest. Further, the analysis outcomes of Western blot showed that anlotinib could down-regulate the expression of Survivin, cyclin D1 and CDK4, as shown in figures 5A and 5B. Collectively, these data suggested that anlotinib sensitized HCT-8/5-FU cells by reducing the cell cycle at S phase.

**Figure 2.**
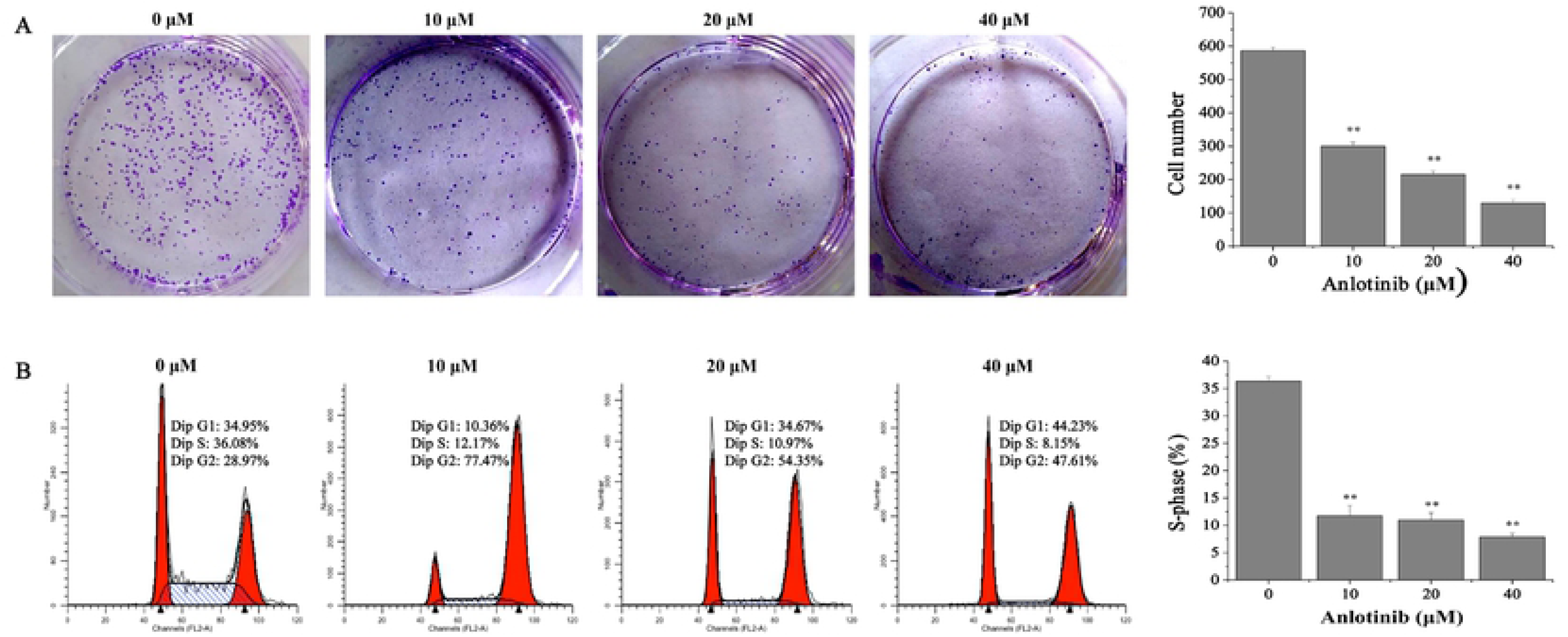
Anlotinib inhibited the proliferation of HCT-8/5-FU. Compared to untreated controls, HCT-8/5-FU were cultured with different concentrations (0, 10, 20, 40 µM) of anlotinib for 24 h, and colony formation and cell cycle was analyzed. (A) Anlotinib decreased the cell number in s dose-dependent manner. (B) Anlotinib arrested the cell cycle at G0/G1 phase in 5-FU-resistant colorectal cancer cells. **P<0.01.

**Figure 3.**
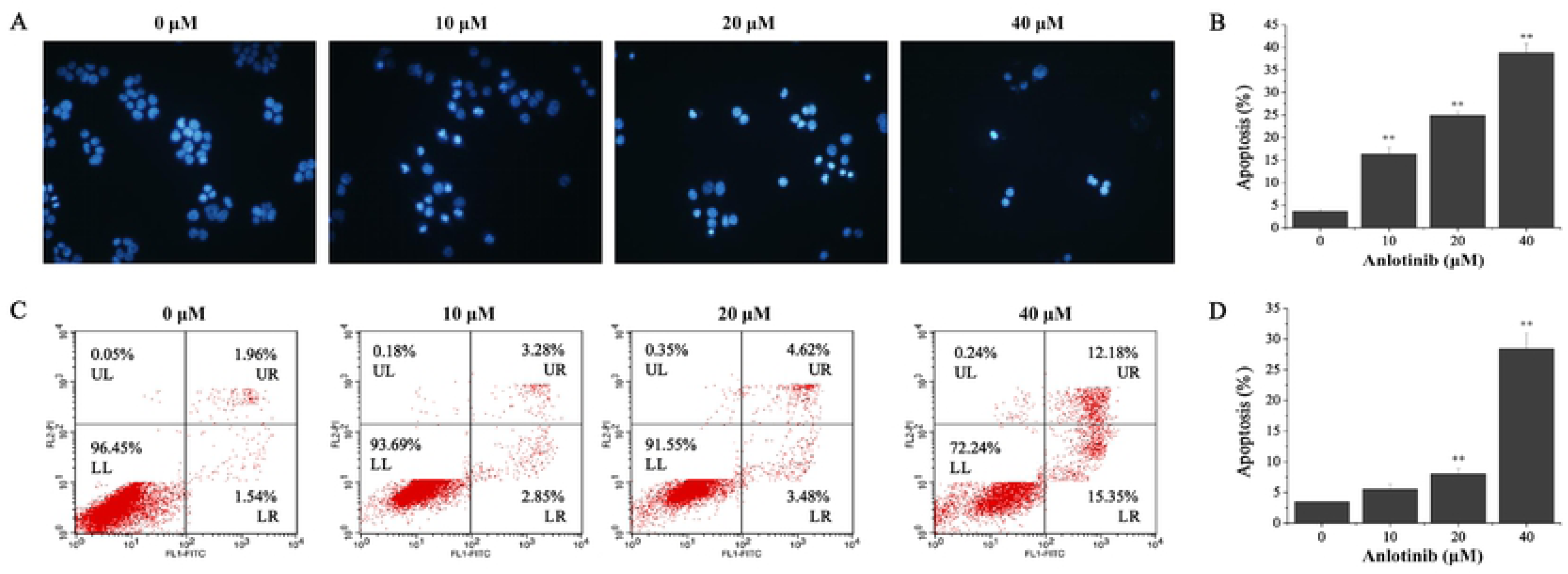
Anlotinib accelerated apoptosis in HCT-8/5FU cells. (A) DAPI staining was used to evaluate apoptotic cells treated with different concentrations. (B) HCT-8/5FU cells were cultured with anlotinib (10, 20, 40 µM) for 24 h and the process of apoptosis detected by flow cytometry. The ratio of apoptotic cells and DAPI are exhibited. **P<0.01.

**Figure 4.**
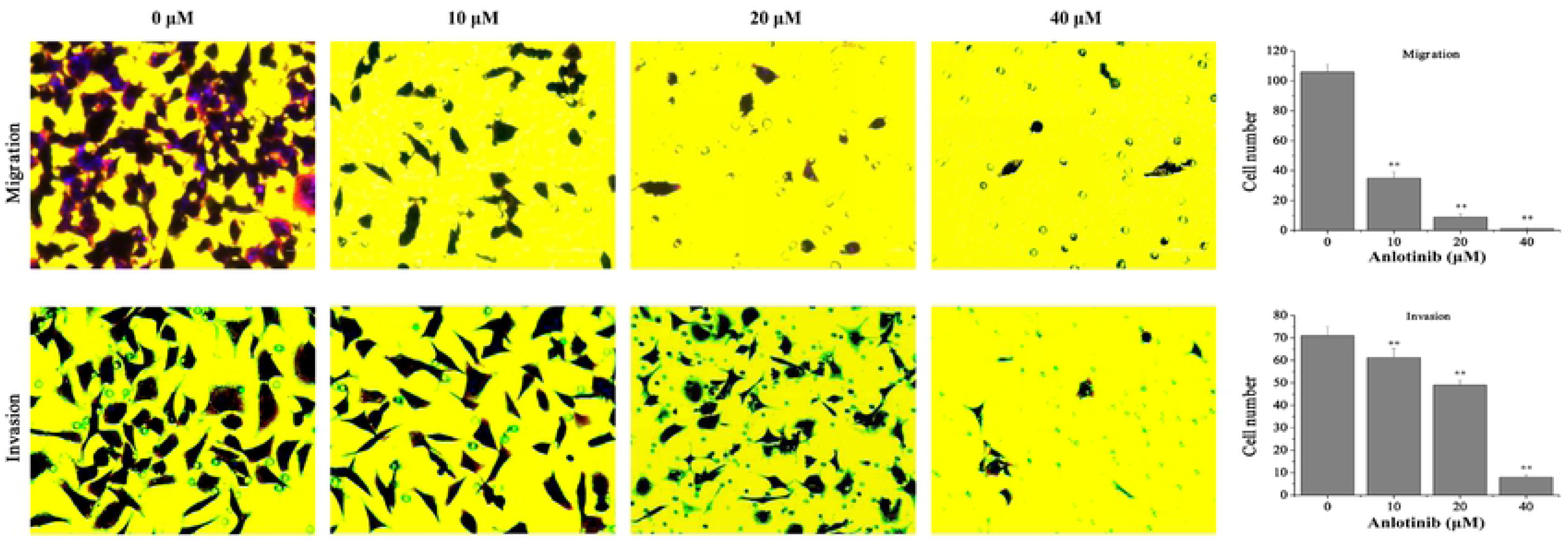
Anlotinib inhibited the migration and invasion of HCT-8/5-FU. HCT-8/5-FU cells were cultured with anlotinib (0, 10, 20, 40 µM) for 24 h. (A) Migration was determined; (B) Invasion assay was carried out. **P<0.01.

**Figure 5.**
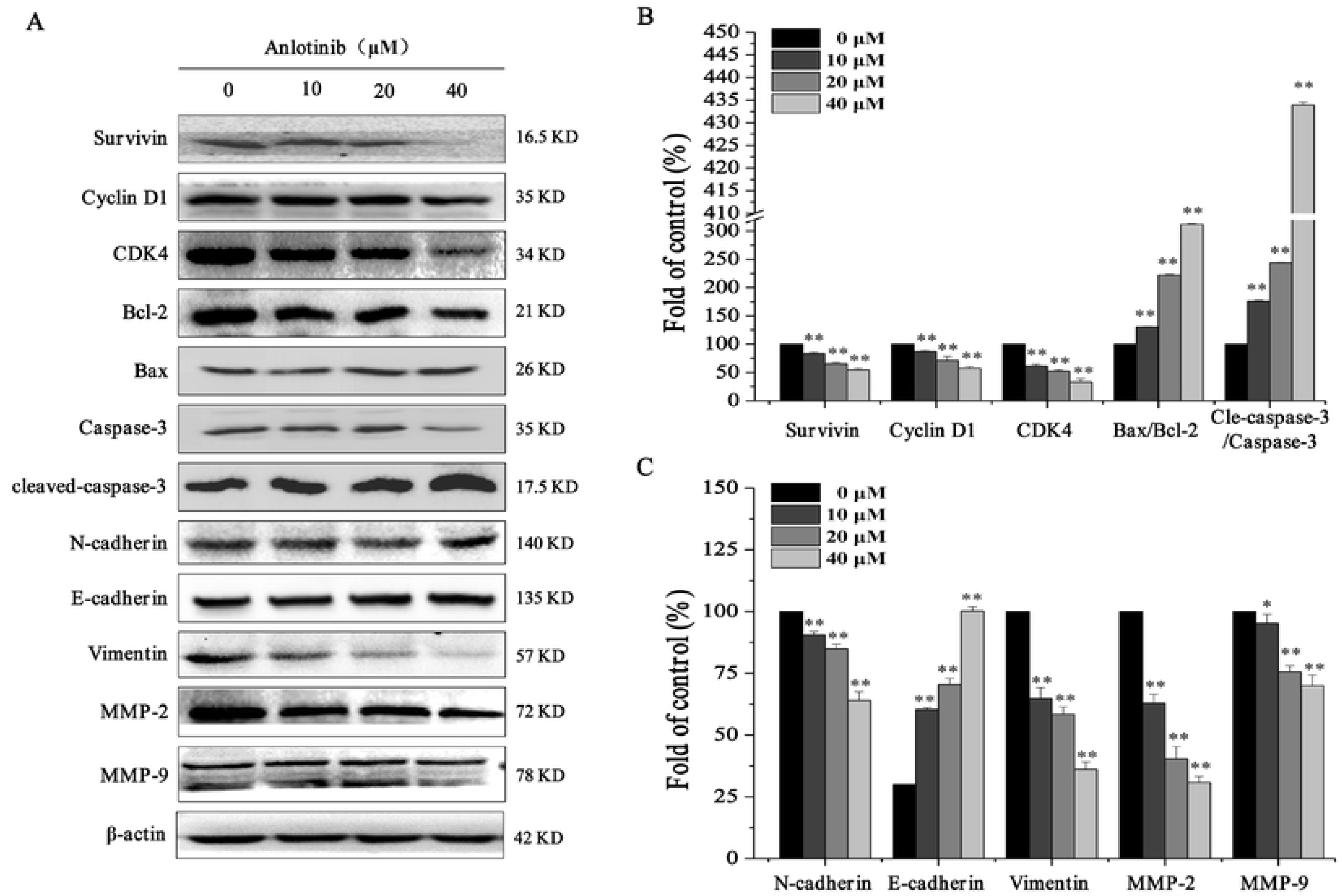
Anlotinib regulated protein expression. HCT-8/5FU cells were cultured with anlotinib (0, 10, 20, 40 µM) for 24 h. Then correlated protein cells were collected for Western blot and analyzed. (A) Expression of protein. (B) Quantitative analysis of cyclin D1, CDK4, survivin, Bax/Bcl-2 and cleaved-caspase-3/caspase-3. (C) Quantitative analysis of E-cadherin, N-cadherin, vimentin, MMP-2 and MMP-9. *P<0.05, **P<0.01.

### Anlotinib induced apoptosis of HCT-8/5-FU cells

In order to confirm whether anlotinib inhibited cell growth by causing apoptosis, we tested its pro-apoptotic activity in HCT-8/5-FU cells via DAPI and Annexin-V/PI staining. Compared to control cells, anlotinib decreased the cell numbers of HCT-8/5-FU and induced dot-like apoptotic body formation in HCT-8/5FU cells in a dosage-dependent pattern, as shown in Figure 3A. To check the pro-apoptosis feature of anlotinib, Annexin V/PI staining was demonstrated by flow cytometry. As shown in Figure 3B, cells were treated with varying concentrations (0, 10, 20 and 40 μM) of anlotinib, and the percentage of apoptotic cells were 3.47 ± 0.03%; 5.54 ± 0.77%; 8.02 ± 0.80% and 28.44 ± 2.54%, respectively. Moreover, expression of protein-related apoptosis as Bcl-2, Bax, caspase-3 and cleaved caspase-3 were tested. As shown in figures 5A and 5B, treatment with 0, 10, 20 and 40 μM of anlotinib up-regulated the expression of Bax and cleaved caspase-3, but inhibited Bcl-2 and caspase-3. Together, these data indicated that anlotinib’s ability to sensitize HCT-8/5-FU may be due to its pro-apoptotic activity.

### Anlotinib inhibited migration and invasion of HCT-8/5-FU cells

To detect the effect of anlotinib on metastasis of HCT-8/5-FU cells, migration and invasion tests were carried out. The experiments showed anlotinib could inhibit the migration and invasion of HCT-8/5-FU cells in a dosage-dependent fashion. As shown in Figures 4A and 4B, treatment with 0, 10, 20 and 40 μM of anlotinib decreased the migratory cells by 106 ± 5, 35 ± 4, 9 ± 2, and 1 ± 1, respectively, and invasive cells by 71 ± 4, 61 ± 4, 49 ± 2 and 8 ± 1, respectively. To investigate the inherent mechanism of the inhibitory activity of anlotinib against endothelial to mesenchymal transition (EMT) in greater depth, the effect of anlotinib on related proteins was investigated. As shown in Figures 5A and 5C, anlotinib was observed to significantly decrease the expression levels of N-cadherin, MMP-2, MMP-9 and vimentin, but increase the expression of E-cadherin.

### Anlotinib inhibited the activation of the PI3K/AKT pathway in HCT-8/5-FU cells

Phosphatidylinositol 3-kinase/protein kinase B (PI3K/Akt) is a common cancer-related signaling pathway associated with cell proliferation, apoptosis, survival, migration, invasion, metastasis and other intracellular transport functions during the origination and development of tumor cells[12, 13]. To clarify the mechanism of action of anlotinib on the PI3K/AKT pathway, we determined the expression of PI3K, AKT and p-AKT, results as in figure 6A and 6B showing anlotinib significantly inhibited the PI3K/AKT pathway. Next, LY294002 (an inhibitor of PI3K) was used. We found that LY294002 could block the activity of PI3K and p-AKT, as shown in Figures 6C and 6D. These data indicated that the reason anlotinib was able to sensitize HCT-8/5-FU cells that it could block the PI3K/AKT pathway.

**Figure 6.**
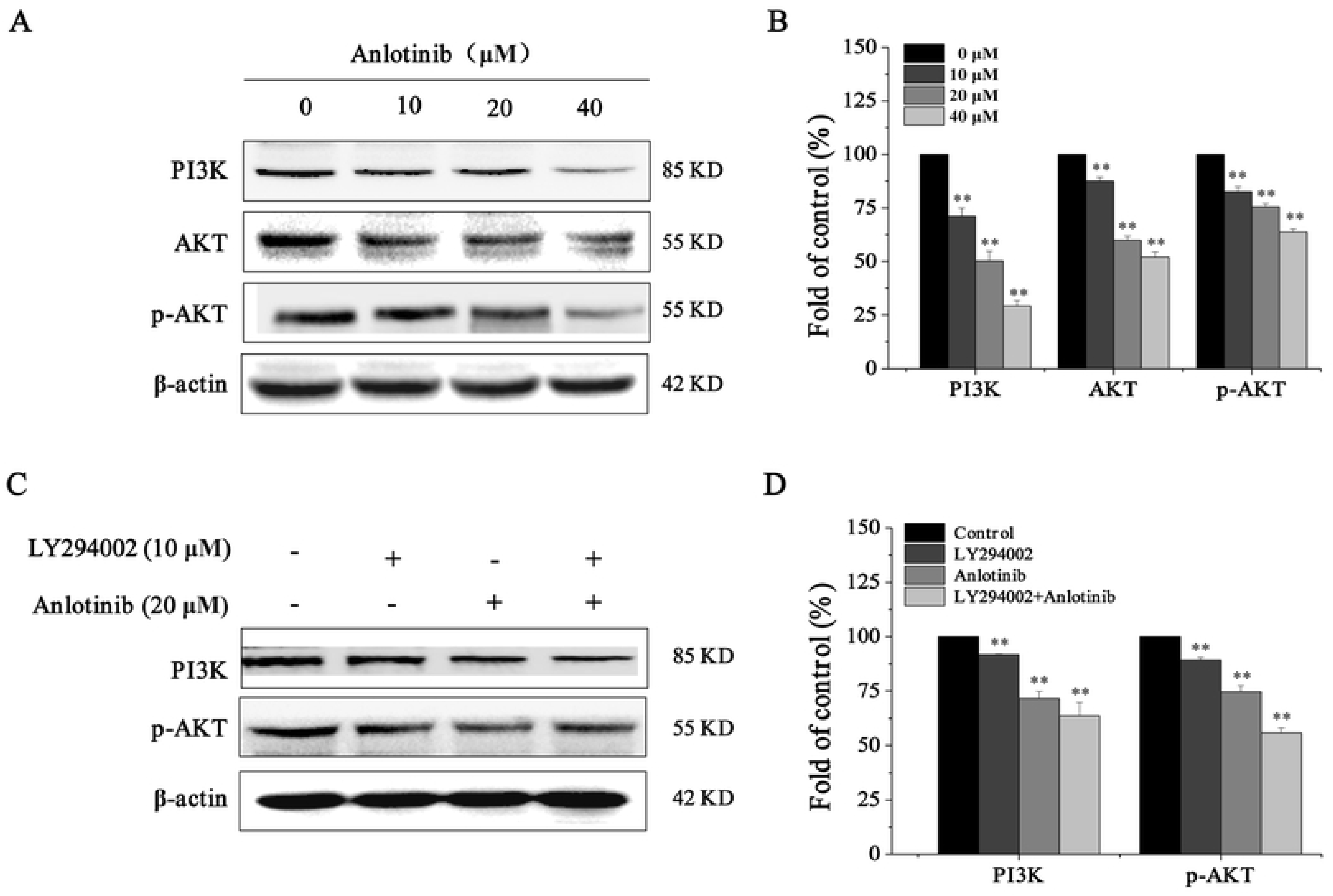
Anlotinib blocked the activity of PI3K/AKT pathway. HCT-8/5FU cells were cultured with anlotinib or LY294002 for 24 h. The proteins of cells were harvested for Western blot. (A) Western blot analysis was used to measure PI3K/AKT related proteins, including PI3K, AKT and P-AKT. (B) Relative quantification of each protein is displayed. (C) LY294002 (the inhibitor of PI3K) was used to block the PI3K/AKT pathway. (D) Relative quantification of the protein. **P<0.01.

## Discussion

Despite recent advances in treatment options, colorectal cancer has remained a deadly disease for many years, with an increasing global incidence[14, 15]. It is widely known that drug resistance, recurrence and metastasis of tumor cells remain the main reasons for chemotherapy failure. The multidrug resistance of tumor cells make chemotherapy drugs lose their efficacy, which increases the difficulty of tumor treatment and reduces the survival rate of tumor patients[16]. Recently, molecular-targeted agents[17], such as cetuximab and panizumab[18], bevacizumab[19, 20], regofenib[21], remoluzumab[22, 23] and RTKs[24, 25] have been used clinically in the treatment of CRC. However, little research has been done on multidrug-resistant cells in colorectal cancer.

Anlotinib is a new type of oral small molecule, multi-target tyrosine kinase inhibitor (TKI). On May 9, 2018, the State Food and Drug Administration (CFDA) officially approved the third line therapy program for terminal non-small-cell carcinoma (NSCLC) patients with anlotinib hydrochloride[26, 27]. Anlotinib had approved anti-tumor for several cancers, such as AMTC, MRCC, NSCLC, and ASTS[28, 29]. Currently, most researches focus on the anti-angiogenic effect of drugs on tumor cells and the therapeutic effect on non-small cell carcinoma[9, 10, 26, 27]. However, relevant data indicated that the effect of anlotinib on the proliferation and apoptosis of drug-resistant cells in colorectal cancer is still rarely reported. Therefore, we believe that the study of anlotinib hydrochloride has great significance for multi-drug-resistant tumor cells.

It is well established that many cell proliferation, survival, adhesion and migration functions are achieved through PI3K/AKT/ERK(MAPK) signaling pathways[30-34]. The pathway of PI3K/AKT cell signaling has significant importance for regulating CRC cell multiplication and EMT[35-38]. After activation, AKT, as a key protein in the pathway, activates or inhibits its downstream target protein through phosphorylation, playing a key part in adjusting cell growth and proliferative effect as well as inhibiting apoptosis[39-41]. In addition, studies have reported that the activation of AKT kinase is essential to a number of events in the pathway of metastasis, containing, tumor microenvironment, tumor immune escape, activation of proliferation, blocking of apoptosis, and activation of angiogenesis[42].

With this knowledge in mind, we studied proliferation and apoptosis, considering that the pathway of PI3K/AKT signaling would become more sensitive in the process of migration and invasion of CRC cells. Hence, we investigated how anlotinib might regulate invasion and migration of HCT-8/5FU cells, with PI3K/Akt signaling pathway continuing to participate in EMT protein expression, so that we might determine the underlying signaling mechanisms. In order to further verify, we also detected cell protein expression and signal transduction pathway activation in CRC through Western blot. Results were that anlotinib hydrochloride could down-regulate the expression of survivin, cyclin D1, CDK4, Bcl-2, caspase-3, N-cad, MMP-2, MMP-9 vimentin and up-regulate the expression of proteins Bax, cleaved caspase-3 and E-cad, by inhibiting the activation of the PI3K/AKT pathway (Figure 6A).

In order to further verify this pathway, we also used PI3K drug inhibitors and measured the changes of PI3K and p-AKT after the addition of inhibitors. The results showed that PI3K inhibitor could reduce the activation of PI3K and p-AKT and that the addition of anlotinib hydrochloride could enhance the inhibitory effect of PI3K and p-AKT (Figure 6B).

As a multiple-target drug, the potential use of anlotinib certainly warrants further investigation, as it may play an important role in other pathways or molecules as well. Although anlotinib is regarded as a promising prospect agent, further in vivo experiments will be needed to profile its side-effects and adverse reactions before its safe clinical application in patients with colorectal cancer can be assured and widely adopted.

## Conclusion

Anlotinib inhibited the proliferation, metastasis and induced apoptosis of HCT-8 / 5-FU cells; and the mechanism could be that anlotinib overcomes multiple drug resistant of the colorectal cancer cells via inactivating PI3K/AKT pathway.

## Data Availability

The data used to support the findings of this study are original and available from the corresponding author upon request.

